# A dynamic 1/f noise protocol to assess visual attention without biasing perceptual processing

**DOI:** 10.1101/2021.07.10.451930

**Authors:** Nina M. Hanning, Heiner Deubel

**Author notes:** corresponding author: Nina M. Hanning.

## Abstract

Psychophysical paradigms measure visual attention via localized test items to which observers must react or whose features have to be discriminated. These items, however, potentially interfere with the intended measurement as they bias observers’ spatial and temporal attention to their location and presentation time. Furthermore, visual sensitivity for conventional test items naturally decreases with retinal eccentricity, which prevents direct comparison of central and peripheral attention assessments. We developed a stimulus that overcomes these limitations. A brief oriented discrimination signal is seamlessly embedded into a continuously changing 1/f noise field, such that observers cannot anticipate potential test locations or times. Using our new protocol, we demonstrate that local orientation discrimination accuracy for 1/f filtered signals is largely independent of retinal eccentricity. Moreover, we show that items present in the visual field indeed shape the distribution of visual attention, suggesting that classical studies investigating the spatiotemporal dynamics of visual attention via localized test items may have obtained a biased measure. We recommend our protocol as an efficient method to evaluate the behavioral and neurophysiological correlates of attentional orienting across space and time.

**Significance statement:** Where (and when) we pay attention can be experimentally quantified via visual sensitivity: Attending to a certain visual signal results in better detection and feature discrimination performance. This approach is widely used, but poses an unrecognized dilemma: The test signal itself, typically a grating or letter stimulus, biases observers’ perception and expectations – and thus also the attention measurement. We developed a stimulus that manages without test items. The signal to measure attention is seamlessly embedded in a dynamic 1/f noise field, so that neither spatial nor temporal information about signal presentation is conveyed. Unlike with conventional approaches, perception and expectations in this new protocol remain unbiased, and the undistorted spatial and temporal spread of visual attention can be measured.

## Introduction

Only a fraction of the information flooding the visual system every time we open our eyes can be processed. Visual attention allows us to selectively focus on specific locations or features while ignoring other aspects of the available information by biasing the neuronal representation of the visual scene (1,2). Depending on the attentional state of the observer, the same retinal input elicits different neurophysiological responses (3-5). As a behavioral consequence of these modulations, attention increases spatial resolution (6,7), enhances contrast sensitivity (8-10), and even alters visual appearance (11,12). Given that stimuli presented in the focus of attention are recognized faster than those appearing outside the focus (13), the spatial deployment of attention is frequently deduced from manual response times. Reaction times, however, reflect the combined effect of detection-, decision- and response-dependent processes (14), which can only be differentiated with special methods (e.g., speed–accuracy trade-off) and models (e.g., drift-diffusion model).

A more direct behavioral correlate of visual attention can be obtained by measuring the sensitivity to discriminate visual features (15). Studies measuring discrimination performance can isolate the different components by rule out speed-accuracy trade-offs and/or rely on signal detection theory (SDT), which indexes sensitivity and criterion separately. In a typical paradigm, the spatiotemporal dynamics of visual attention are measured by briefly presenting a test stimulus (commonly among several distractors) at a specific point in time and at one out of several pre-specified locations across the visual field. Observers are instructed to discriminate a target-specific feature or its identity. Since attention enhances visual processing, a higher discrimination performance (measured as %-correct responses) or increased visual sensitivity (measured as d-prime) for a particular item reflects the allocation of attention toward its location.

The discrimination approach has become highly popular and a variety of different discrimination features, such as stimulus identity (e.g., 16), orientation angle (e.g., 11), or motion direction (e.g., 17) have been employed (see 18 for a comparison of the most popular stimuli). However, these conventional approaches face a common shortcoming: They all rely on localized test stimuli to determine the deployment of visual attention. The potential problem with this approach is that these stimuli could bias visual perception by structuring the visual field (19-22) and may thus affect what they are intended to measure – the spatial distribution of attention. In a paradigm using a typical stimulus configuration as displayed in **Fig. 1A**, attention is likely biased towards the presented stimuli (as compared to locations in between or further in- or outside), since those are the only locations containing potentially task-relevant visual information (**Fig. 1B**). This bias could occur automatically, in a bottom-up fashion, but might also have a top-down component, in that observers strategically deploy their attention to potential test locations to benefit their task performance. Critically, these biases are not necessarily linear and hard to estimate. They can unsystematically affect different experimental manipulations and thus distort the interferences drawn from the study.

**Figure 1.**
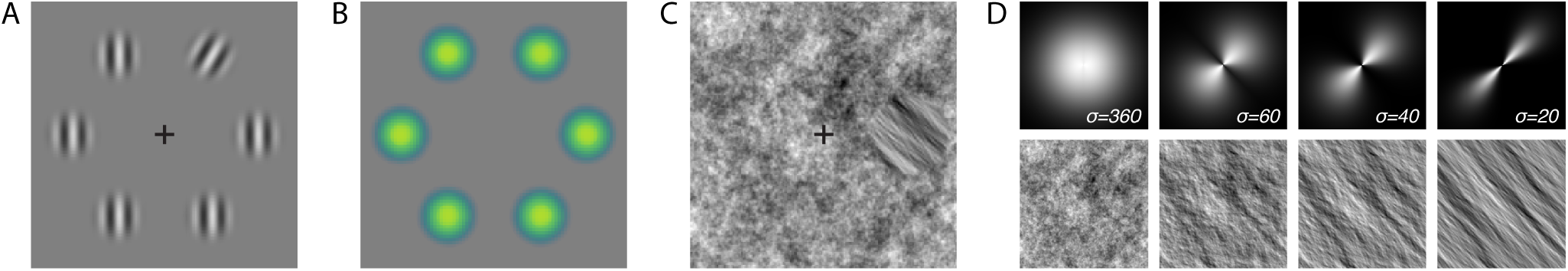
Rational of the dynamic 1/f noise protocol. (**A**) Example of a typical paradigm to investigate visual attention (adapted from 23). Observers discriminate a test stimulus (a clockwise or counterclockwise oriented Gabor) presented along with several distractors (here vertical Gabors). (**B**) The stimulus configuration shown in (A) reveals potential test locations, to which observer’s attention might be biased (visualized by colored blobs). (**C**) The dynamic 1/f noise protocol. Observers discriminate the tilt angle of an orientation filtered noise patch embedded in an unfiltered 1/f noise background (here at 3 o’clock, tilted counterclockwise relative to vertical). The orientated patch can appear at any location in the noise field, potential test locations remain unknown to the observer. (**D**) Examples of orientation filters with different widths (σ; upper row) and the resulting filtered noise image (bottom row). By varying orientation filter width, the difficulty of the discrimination task can be continuously adjusted (the narrower the width, the stronger the signal and the easier the task).

To overcome such attentional biases induced by the experimental design, we developed a dynamic, item-free 1/f (“pink”) noise protocol that allows to assess visual performance across the scene without object-like visual structures. By embedding an orientation discrimination signal into full-field 1/f noise (**Fig. 1C**) observers’ visual sensitivity can be probed at any location in the noise field without revealing a specified set of potential test locations. The local discrimination signal composed of orientation-filtered 1/f noise (displaying, for example, a clockwise or counterclockwise orientation relative to vertical) is inserted into the 1/f noise field with a soft boundary and can take any shape and size. By altering the orientation filter width, the strength of the discrimination signal can be continuously manipulated (**Fig. 1D**), which allows to precisely adjust the overall difficulty of the perceptual discrimination task. Since sudden local changes in the stimulus display capture attention (e.g., 24,25) and potentially bias the temporal dynamics of attention allocation (18), the full-field noise stimulus (including the embedded orientation signal) continuously changes over time. This dynamic approach also permits to present the orientation signal at any time for a desired presentation duration without providing observers with temporal information regarding test signal occurrence. See **Methods** for reference to detailed description and example code.

We selected orientation-filtered 1/f spatial noise as discrimination signal for two major reasons. First, the 1/f falloff of the amplitude spectrum is “naturalistic” in the sense that real-world scenes and textures typically show a similar spatial frequency dependence (26,27). Second – and a decisive advantage of the pink noise stimulus over conventional approaches – this particular type of test signal should be scale-invariant and yield discrimination thresholds that are constant across retinal eccentricities. Spatial acuity decays with increasing distance from the fovea (the center of gaze), i.e., the visual system becomes less sensitive to higher spatial frequencies further in periphery (28,29). As argued in a seminal study by Field (26), early visual processing can be approximated by assuming an array of spatial bandpass filters (spatial frequency channels) representing the response properties of cortical cells, in which the spatial frequency bandwidth is constant in octaves (e.g., 1 to 2 c/deg, 2 to 4 c/deg, etc.). Under this assumption, a 1/f amplitude falloff will yield equal energy in equal octaves (e.g., the energy between 1 and 2 c/deg will be equal to the energy between 2 and 4 c/deg, etc., for a detailed analysis see 26). Since peripheral vision can be well modeled by eccentricity (cortical magnification / M-) scaling (30), it follows that relative energy in the spatial frequency channels is also eccentricity-invariant. As we use orientation filtering that is homogeneous across all spatial frequencies (i.e., the orientation signal contains the full 1/f frequency spectrum), perceptual orientation discrimination thresholds should be largely independent of the eccentricity at which the test signal is presented.

We conducted two experiments to demonstrate the applicability and effectiveness of the dynamic 1/f noise protocol. In Experiment 1 we establish that (1) orientation discrimination accuracy continuously scales with the width of the orientation filter, allowing to adjust discrimination task difficulty (via altering orientation filter width) to individual observers’ capabilities; (2) the embedded orientation signal itself does not capture attention, as it is not discriminable when presented outside the focus of attention; (3) local discrimination accuracy is, as predicted, largely independent of the visual eccentricity at which the test signal is presented, enabling direct comparison of visual performance from central to peripheral locations. In Experiment 2 we demonstrate that (4) items in fact structure the visual field and markedly bias visual perception. This emphasizes the relevance of the item-free 1/f noise protocol and its unique capability of measuring the unbiased distribution of visual attention across time and space.

## Methods

### 1/f noise protocol

The noise stimulus comprises a dynamically changing 1/f noise background, gradually changing from one noise image to another, in which a local orientation signal of desired shape and size is neatly embedded for a desired duration. A detailed description as well as custom MATLAB code for generating the stimulus will be available on GitHub upon manuscript publication.

### Empirical data and experimental procedure

#### Observers

Sample sizes were determined based on previous work (18,31,32). 6 observers (ages 19 - 28 years, 4 female) completed Experiment 1a and Experiment 1b, 6 observers (ages 20 - 31 years, 5 female) completed Experiment 2. All observers were healthy, had normal vision, and except for one author (N.M.H. participated in Experiment 2) were naive as to the purpose of the experiments. The protocols for the study were approved by the ethical review board of the Faculty of Psychology and Education of the Ludwig-Maximilians-Universität München (approval number 13_b_2015), in accordance with German regulations and the Declaration of Helsinki. All observers gave written informed consent.

#### Apparatus

Gaze position of the dominant eye was recorded using a SR Research EyeLink 1000 Desktop Mount eye tracker (Osgoode, Ontario, Canada) at a sampling rate of 1 kHz. Manual responses were recorded via a standard keyboard. The experimental software was implemented in MATLAB (MathWorks, Natick, MA), using the Psychophysics (33,34) and EyeLink toolboxes (35). Observers sat in a dimly illuminated room with their head positioned on a chin-forehead rest. Stimuli were presented at a viewing distance of 60 cm on a 21-in. gamma-linearized SONY GDM-F500R CRT screen (Tokyo, Japan) with a spatial resolution of 1,024 by 768 pixels and a vertical refresh rate of 120 Hz.

#### Experimental design

**Experiment 1a** (see **Fig. 2A**). Each trial began with observers fixating a central black (∼0 cd/m2) fixation dot (radius 0.1°) on gray background (∼60 cd/m2). Once stable fixation was detected within a 1.75° radius virtual circle centered on fixation for at least 200 ms, the trial started with the presentation of circular dynamic 1/f noise background (average luminance ∼60 cd/m^2^), extending across the screen height and windowed by a symmetrical raised cosine (radius 14.32°, sigma 2.39°). The background noise was updated at 60 Hz, changing gradually from one noise image to another within 4 frames. After a random fixation period between 400 and 800 ms, in *cued trials* (half of trials, randomly intermixed) a pink color cue appeared for 75 ms, indicating the location of the upcoming discrimination signal (RGB: [204, 0,102], ∼60 cd/m2, same dimensions as the upcoming discrimination signal). Following a delay of 100 ms, a local orientation signal, oriented 40° clockwise or counterclockwise from vertical, windowed by a symmetrical raised cosine (radius 1.75°, sigma 0.875°), was embedded in the background noise at a randomly selected position at one out of four retinal eccentricities (0°, 3.5°, 7°, or 10.5°). The orientation filter strength *σ* was randomly selected for each trial (9 linear *σ*-steps between 30 and 70). In the *uncued* trials, no color cue was presented and the discrimination signal occurred at an unpredictable location (same eccentricities, timing, and specifications as in cued trials). After 50 ms, the orientation signal was masked by the reappearance of non-oriented noise for 500 ms, before the background noise disappeared, and observers reported the perceived orientation via button press (two-alternative forced choice). They were informed that their orientation report was non-speeded and they received auditory feedback for incorrect responses.

**Figure 2.**
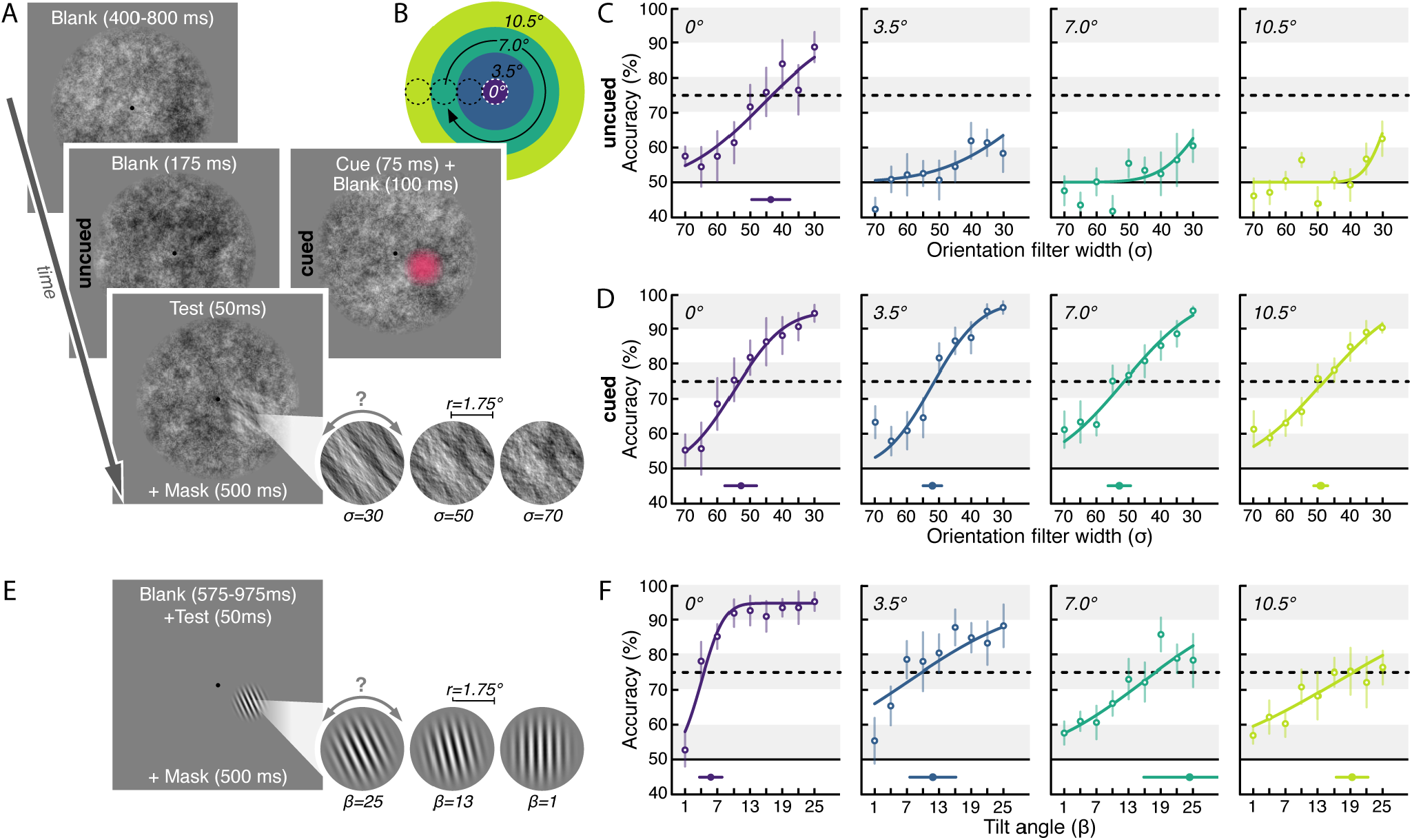
Experiment 1. (**A**) Experiment 1a. Observers (*N* = 6) fixated a central fixation dot on dynamic 1/f noise background and aimed to discriminate a local orientation signal. This signal was presented at different eccentricities (see B) and either was preceded by a 100% valid color pre-cue (*cued*; results in D), or not (*uncued*; results in C). (**B**) Visualization of the tested eccentricities. Color code matches the psychometric functions in C, D, and F. (**C&D**) Group-averaged psychometric functions (discrimination performance as a function of orientation filter width) for *uncued* (C) and *cued* trials (D). 75% discrimination thresholds (Th^75^) derived from individual observers’ psychometric functions are shown at the bottom of each plot with filled dots. Error bars denote the SEM. (**E**) Experiment 1b. The same observers fixated a central fixation dot on uniform gray background and aimed to discriminate the orientation of a Gabor. The Gabor had the same size as the orientation signal in Experiment 1a and was presented at same eccentricities (see B). (**F**) Group-averaged psychometric functions for Experiment 1b. Conventions as in C&D.

**Experiment 1b** (see **Fig. 2E**). Task and timing were identical to Experiment 1a with the following differences: No dynamic 1/f noise background was presented. Instead, upon trial start a local stimulus stream alternating at 20 Hz between a vertical Gabor patch (2 cpd, random phase, full contrast, ∼60 cd/m2) and a Gaussian pixel noise mask (composed of ∼0.25°-wide pixels ranging randomly from black to white, ∼60 cd/m2) was presented (18,42). The stream had the same size / raised cosine envelope (radius 1.75°, sigma 0.875°) and was presented at the same retinal eccentricities as the 1/f orientation signal in Experiment 1a (0°, 3.5°, 7°, or 10.5°). For a duration of 50 ms, the stream contained an orientation signal – a Gabor patch rotated clockwise or counterclockwise relative to the vertical. The tilt angle β was randomly selected for each trial (9 linear β-steps between 25° and 1°). To avoid apparent motion effects, after orientation signal presentation the stream continued with alternating noise patches and blanks. Observers indicated via button press in a non-speeded manner whether they had perceived the orientation to be tilted clockwise or counterclockwise, and received auditory feedback for incorrect responses.

Observers performed 15 experimental blocks (10 of Experiment 1a, 5 of Experiment 1b), each of 180 trials. Trials in which we detected blinks, accidental eye movements, or broken eye fixation (gaze deviating further than 1.75° from the instructed fixation location) were discarded. In total we included 10,223 trials in the analysis of the behavioral results for Experiment 1a (1,704 ± 81 trials per observer; mean ± SEM) and 5,328 trials for Experiment 1b (888 ± 3 trials per observer).

**Experiment 2** (see **Fig. 3A**). On a rectangular dynamic 1/f noise field (height 2.5° x width 20°, average luminance ∼60 cd/m^2^) we presented five black (∼60 cd/m^2^) circular frames (radius 1.0°), evenly spaced on the horizontal midline (± 0°, 4°, and 8° relative to noise background center). Observers were required to fixate the center of the middle frame. Once stable fixation was detected within a 1.75° radius around the fixation location for at least 200 ms, the trial started with a 300 - 1000 ms fixation period (duration randomly chosen), after which we embedded a local orientation signal (tilted ± 40° relative to vertical) in the background noise, windowed by a symmetrical raised cosine (radius 1.0°, sigma 0.875°). The orientation signal was presented at one out of 9 evenly spaced horizontal positions (± 0°, 2°, 4°, 6°, and 8° relative to central fixation), thus either centered within one of the frames or in between them. Observers were informed that the orientation signal would occur at each of the 9 test locations with equal probability. After ∼42 ms, the orientation signal was masked by the reappearance of non-oriented noise. Following a masking period of 300 - 650 ms, the dynamic 1/f background noise disappeared and observers indicated via button press in a non-speeded manner whether they had perceived the orientation to be tilted clockwise or counterclockwise.

**Figure 3.**
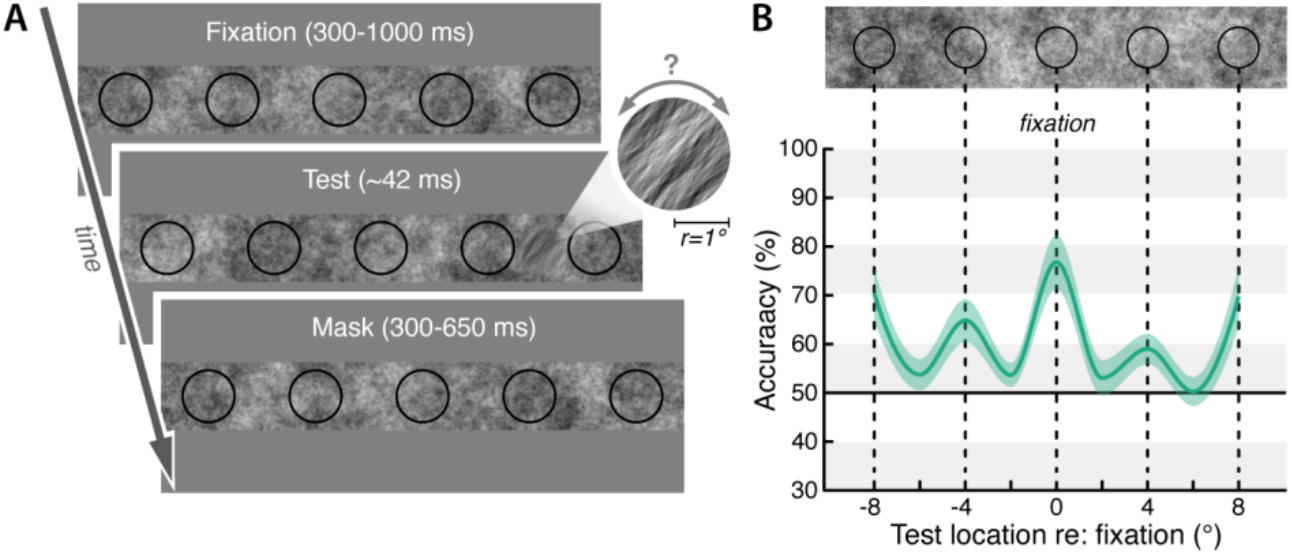
Experiment 2. (**A**) Experimental design. Observers (*N* = 6) fixated the central of five black frames presented on dynamic 1/f noise background. A brief orientation signal (here clockwise) was embedded either within or in between the frames. (**B**) Group average orientation discrimination performance (green line) as a function of horizontal test signal position (corresponding stimulus arrangement shown on top). The colored area indicates the SEM.

To ensure a comparable performance level across individual observers, the orientation signal strength was adjusted via the width of the orientation filter in a threshold task preceding the experiment. The experimental design resembled the main experiment, but no circular frames were presented and the orientation signal was preceded by a 100% valid spatial pre-cue (same cue as in Experiment 1a). For each trial we randomly selected the orientation filter strength out of 5 linearly spaced filter widths (*σ* 10 to 50; note that we used narrower filter values compared to Exp. 1a due to the shorter presentation time and smaller size of the orientation signal). We determined the filter width corresponding to 90% discrimination accuracy individually for each observer by fitting cumulative Gaussian functions to the average discrimination performance per filter width σ (collapsed across signal locations) via maximum likelihood estimation, and used the determined filter width for the main experiment.

Observers performed 150 trials of the threshold task and 347 - 415 trials of the main experiment. Trials in which we detected blinks, accidental eye movements, or broken eye fixation (gaze deviating further than 1.75° from the required fixation position) were discarded. In total we included 2,289 trials in the analysis of the behavioral results for Experiment 2 (382 ± 11 trials per observer).

#### Behavioral data analysis and visualization

For **Experiment 1**, we fitted cumulative Gaussian functions to each observer’s average discrimination accuracy (% correct) per filter width *σ* (Exp. 1a, **Fig. 2C&D**; also Exp. 2 - threshold task) or tilt angle *β* (Exp. 1b, **Fig. 2F**), separately for each test eccentricity. Based on these psychometric functions, we estimated each observer’s 75% discrimination threshold (Th^75^) and plotted the averaged estimates (group-mean ± SEM) together with a group-averaged psychometric function. To visualize discrimination accuracy across space in **Experiment 2**, we interpolated between the group-averaged discrimination accuracy (% correct) for each test location (± 0°, 2°, 4°, 6°, and 8° relative to central fixation).

For statistical comparisons we used permutation tests to determine significant performance differences between two conditions or locations. We resampled our data to create a permutation distribution by randomly rearranging the labels of the respective conditions for each observer and computed the difference in sample means for 1000 permutation resamples (iterations). We then derived *p*-values by locating the actually observed difference (difference between the group-averages of the two conditions) on this permutation distribution, i.e. the *p*-value corresponds to the proportion of the difference in sample means that fell below or above the actually observed difference (*p*-values were Bonferroni-corrected for multiple comparisons).

## Results

To demonstrate that the noise stimulus can accurately measure visual sensitivity across the scene, in **Experiment 1a** we tested local 1/f noise orientation discrimination performance at various retinal eccentricities with and without spatial attention focused on the test signal location. Observers fixated a central fixation target presented on a circular, dynamic 1/f noise background (**Fig. 2A**). After a fixation period, a local orientation signal was presented for 50 ms, either centered on the fixation or at a randomly chosen position 3.5°, 7°, or 10.5° (degree of visual angle) away from fixation (**Fig. 2B**). In half of the trials (*cued trials*; see **Supplemental Video S1**), we manipulated the allocation of spatial attention by presenting a brief, salient color pre-cue that indicated the test signal location shortly before signal onset (100% valid); in the other half or trials (*uncued trials*; see **Supplemental Video S2**) no information regarding signal location or presentation time was provided. At the end of each trial, observers indicated via button press whether they had perceived an orientation clockwise or counterclockwise relative to vertical (two-alternative forced choice task).

When attention was not directed via the spatial pre-cue (*uncued trials*; **Fig. 2C**), discrimination performance scaled with the width of the orientation filter (used to create the orientation signal) if the orientation signal was presented at the center of gaze (0° eccentricity from fixation): the narrower the filter (i.e., the stronger the orientation signal), the higher the proportion of correct orientation judgements. However, for all peripheral test locations (3.5°, 7.0°, and 10.5° eccentricity), observers’ accuracy was low and hardly increased with decreasing filter width. Thus, when observers were unaware of test location and presentation time, they were able to successfully discriminate the orientation signal only when it occurred right at the center of their gaze.

The superior discrimination performance at the central compared to the peripheral locations may not seem surprising, considering that the visual system’s acuity and perceptual sensitivity is highest at the fovea and decays towards the periphery (28,29). As a consequence, we clearly perceive fine details in the center of gaze (where we are sensitive to high spatial frequencies), that we cannot distinguish when viewing them peripherally (where we are more sensitive to lower spatial frequencies). The 1/f property of the orientation signal, however, should allow equal orientation discriminability across eccentricities. This was in fact the case when the 1/f orientation signal was preceded by a salient, 100% valid pre-cue, which can be assumed to automatically attract exogenous spatial attention to its location (2,29). Discrimination accuracy in the *cued trials* (**Fig. 2D**) increased with decreasing filter width in a highly similar fashion for all tested eccentricities. The derived 75% perceptual discrimination thresholds (see **Methods**) remained stable from the fovea (Th^75^ 0°: *σ =* 52.84 ± 4.83; mean ± SEM) up to the furthest tested retinal eccentricity (Th^75^ 3.5°: *σ =* 52.11 ± 2.91, 7.0°: *σ =* 52.91 ± 3.50, 10.5°: *σ =* 49.13 ± 2.08; Th^75^ 0° vs. 10.5°: *p* = 0.304). This indicates that local 1/f orientation discrimination performance is largely independent of test signal eccentricity – as long as the signal is presented within the focus of attention. When not experimentally manipulated, attention is normally coupled to the focus of gaze, which explains why the orientation signal could only be well discriminated at the central fixation location. This location is also where observers should anchor their attention focus to maximize chances of capturing the discrimination signal occurring anywhere in the surrounding noise field. The observation that even strong 1/f orientation signals (created with a narrow filter) could not be discriminated peripherally without spatial attention being deployed towards them (**Fig. 2C**, *uncued trials* 3.5° - 10.5°) demonstrates that the orientation signals do not pop-out, i.e., they do not themselves attract attention.

In summary, we established a consistent relationship between orientation filter width and discrimination accuracy. Discrimination task difficulty thus can be continuously manipulated by adjusting the orientation filter width. Moreover, and of particular relevance for the utility of the noise protocol, local 1/f orientation discrimination performance is largely independent of visual eccentricity. This is a key advantage over conventional psychophysical protocols, for which discrimination performance naturally decreases with increasing distance from fixation.

We illustrate this common issue in **Experiment 1b**, using an experimental design that was matched to Experiment 1a (**Fig. 2E**). The same observers were asked to discriminate the orientation (clockwise vs. counterclockwise) of Gabor patches that had a fixed size, contrast, and spatial frequency and were presented at the different retinal eccentricities used in Experiment 1. Akin to manipulating orientation filter width (Experiment 1a), we varied the tilt angle – a common approach to titrate orientation discrimination task difficulty in a Gabor-protocol. **Fig. 2F** shows the typical eccentricity-dependence seen in conventional perceptual tasks: Foveal discrimination performance sharply increased as a function of tilt angle, yielding a low 75%-threshold (Th^75^ 0°: *β* = 5.80 ± 2.10). Across visual eccentricities, however, orientation discrimination accuracy decreased, requiring larger tilt angles for more eccentric test locations to achieve equivalent 75% discrimination accuracy (Th^75^ 3.5°: *β* = 11.84 ± 4.29, 7.0°: *β* = 24.35 ± 8.54, 10.5°: *β* = 19.30 ± 2.96; Th^75^ 0° vs. 10.5°: *p* = 0.001). This decay of visual sensitivity with increasing distance from the fovea is common for conventual perceptual measures and can be counteracted by up-scaling stimulus size (according to cortical magnification), increasing contrast, or lowering spatial frequency information. Experiment 1a shows that such adjustments are not necessary with the pink noise stimulus, since the 1/f property ensures that the same orientation signal can be discriminated equally well at foveal and peripheral locations.

We made use of this feature in **Experiment 2**, in which we evaluated the protocol’s capability of assessing discrimination performance across the visual field, as well as the potential shaping influence of items on this measurement. We presented five circular frames on a rectangular dynamic 1/f noise field and asked observers to fixate the center of the middle item (**Fig. 3A**). Following a fixation period, we briefly presented a local 1/f orientation signal at one out of nine locations along the horizontal (± 0°, 2°, 4°, 6°, or 8° relative to the center) – either inside one of the frames or between them. After a masking period, observers indicated their orientation discrimination judgement. Critically, the presented frames were completely task-irrelevant, and the critical orientation signal was equally likely to occur at any of the evenly spaced locations (independent of whether the location was framed by an outline or not). To ensure a comparable discrimination performance across observers, orientation signal strength was individually adjusted prior to the experiment (see **Methods**).

As suspected, the presence of the visual outlines selectively modulated perception. Compared to the average discrimination accuracy for orientation signals presented outside the frames (52.53 ± 1.91%; mean ± SEM), local discrimination performance was significantly improved whenever the orientation signal occurred within one of the frames. For each observer we found spatially specific peaks in orientation discrimination accuracy inside frames at all three tested eccentricities (**Fig. 3B**; framed signal at central 0°: 76.67 ± 5.48% vs. no-frame: *p* = 0.003; framed signal at ± 4°: 61.71 ± 2.18% vs. no-frame: *p* = 0.003; framed signal at ± 8°: 69.99 ± 4.27% vs. no-frame: *p* = 0.003). The consistent sensitivity benefit for signals presented inside a visual structure demonstrates that our perception is markedly shaped by the presence of items in the visual scene. Critically, this item-induced bias was uneven across different eccentricities (effect size 0°: *d* = 2.63; 4°: *d* = 2.01; 8°: *d* = 2.36). This demonstrates that visual structures pose the risk of compromising the attention measurement in a non-systematic way. In line with previous work showing the impact of placeholder objects on attentional modulations of visual perception (19-21), these findings underline the necessity of an unbiased, item-free approach to map perceptual dynamics across the visual field in the absence of object-like structures.

## Discussion

Conventional psychophysical paradigms to assess visual attention rely on localized test items, which may have the adverse side effect of distorting the attention measurement by shaping visual perception across the scene. We created a full-field 1/f noise stimulus that allows for an unbiased investigation of attention across space and time by meeting the following criteria: (1) The protocol assesses visual attention without object-like structures, as to not artificially bias attention towards potential test locations. This is achieved by seamlessly integrating the orientation signal into full-field 1/f noise – preventing both automatic bottom-up biases (i.e., prioritized processing of locations containing visual items as compared to “empty” locations), as well as strategic top-down biases (i.e., observers anticipating potential test locations and selectively deploying processing resources towards them). (2) To not convey any information about presentation time and duration of the test stimulus, the orientation signal is temporally embedded into a continuously changing full-field noise background. We achieved this by dynamically updating the 1/f noise field, such that local signal onset and offset are concealed by continuous full-field changes. (3) The overall discrimination task difficulty can be precisely adjusted. In the 1/f noise protocol, task difficulty scales with orientation filter width: a narrower filter yields a stronger, thus better distinguishable orientation signal. (4) Finally, the orientation signal blends sufficiently well into the background pattern, both spatially and temporally, as to not ‘pop-out’ (i.e., attract attention itself), which is the case with conventional, abrupt-onset stimuli (24,25,36,37).

We conducted a first experiment to verify that the 1/f noise protocol in fact meets the criteria established above. **Experiment 1a** demonstrates that discrimination accuracy indeed scales with orientation filter width: When presented in the focus of attention, narrower orientation signals were discriminated with increasingly higher accuracy. This validates that task difficulty can be flexibly calibrated – an essential requirement to prevent floor and ceiling effects (i.e., the discrimination task is too hard / easy) and to account for inter-individual performance differences as well as potential learning effects across multiple test sessions or experiments (18). The relationship between orientation filter width and discrimination performance is reminiscent of the link between contrast strength and visual performance: Increasing stimulus contrast, similar to narrowing the orientation filter, results in enhanced orientation discrimination accuracy (9,38,39). While the effect of stimulus contrast on visual performance has been already linked to neurophysiology (40), future studies need to evaluate a potentially similar relationship between 1/f orientation filter width (and visual performance) and neural responses. Such link, akin to the established contrast response function, would qualify the 1/f noise stimulus to investigate the computational and neurophysiological mechanisms underlying the effects of different types of attention on visual perception.

Our results furthermore show that the orientation signal blends sufficiently well with the background that it can only be successfully discriminated when it is attended: If the signal was not preceded by a salient cue biasing spatial attention to the test signal location, observers’ discrimination performance remained at chance level. This verifies that the orientation signal, as intended, did not ‘pop-out’ – which is a common undesired side effect of transient visual signals (24,25,36,37). Discrimination signals that ‘pop-out’ can be assumed to attract attention and therefore to bias the attention measurement. In conventional paradigms, attentional pop-out is typically avoided by reducing the overall baseline performance via increasing the number of distractor items presented alongside the discrimination target (41-43). Our data demonstrate that the 1/f noise stimulus allows a subtle assessment of attention without employing any distractors. It therefore is a promising tool for investigation spatial and temporal perceptual dynamics, also in the context of motor actions. Discrimination signals that capture attention not only prevent valid conclusions about the actual distribution of attention in space, but also interfere with motor programming: We recently observed increased saccade latencies when sudden-onset test targets were presented during movement preparation (18). This effect is compatible with the phenomenon of *saccadic inhibition*, whereby a transient change in the scene causes a depression in saccadic frequency approximately 100 ms thereafter (44,45). Given the tight coupling of eye movement preparation and visual attention (16,23,46,47), interrupting saccade preparation may likely also affect the temporal dynamics of visual attention. Embedding the discrimination signal in a continuously changing display, such as a dynamic noise background, prevents this potential interference (18).

The evaluation of perceptual thresholds across retinal eccentricities confirms that 1/f orientation signals can be equally well distinguished at foveal and peripheral locations. This is an outstanding property specific to the 1/f characteristic of the noise stimulus. As exemplified in **Experiment 1b**, sensitivity for most visual features naturally decreases with increasing distance from the center of gaze (28,48). Spatiotemporal maps of visual attention created with conventional test stimuli are affected by these perceptual inhomogeneities, which prevents a direct comparison of attentional effects on foveal and peripheral visual performance. To account for this confound, the discrimination signal strength needs to be adjusted separately for each tested retinal eccentricity (e.g., 21,49), which is time consuming, comparably more error-prone, and less flexible. Since local 1/f orientation sensitivity is largely constant across visual eccentricities, the noise stimulus is resistant to visual sensitivity changes across space and thus offers the unique opportunity to directly map attentional modulations of visual perception continuously across space, without the need to increase discrimination signal strength with increasing test eccentricity.

Another key advantage of the noise protocol is its capability of assessing visual sensitivity without actual test items. In **Experiment 2** we demonstrate why this is crucial for an unbiased assessment of visual performance across space: The circular outlines that we presented on the otherwise item-free noise field markedly biased visual perception, as they elicited consistent, spatially specific performance benefits inside each visual structure. This verifies the hypothesized shaping impact of even task-irrelevant items on visual sensitivity across space. It is to be expected that conventional paradigms, that rely on the presentation of discrete test and distractor items likewise bias processing resources towards the presented stimuli (as compared to item-free space). In addition to this automatic, bottom-up component, observers likely strategically prioritize potential target locations, since those may contain task-relevant information. It has been shown that (un)certainty concerning the target stimulus location – experimentally manipulated by indicating the size of the to be attended area (i.e., the “attention field”) with placeholder items – affects the mechanisms by which attention modulates visual perception and the underlying neuronal response (50-52). This demonstrates that the spatial distribution of processing resources across the visual field is markedly shaped by scene-structuring objects – affecting both behavioral and neurophysiological correlates of visual perception – and highlights the relevance of a protocol that allows to measure attention in an item-free, unbiased manner.

Since the noise protocol is not bound to local test items, the orientation signal can take any shape and size, and can be embedded into its background with soft boundaries for any desired presentation duration. This offers a new opportunity to directly investigate the temporal and spatial properties of the focus of attention and reconcile previous findings based on which it has been characterized as a moving spotlight (e.g., 53,54), as a zoom lens (e.g., 55), a gradient of processing resources (e.g., 56), or a Mexican hat (e.g., 57,58). The noise protocol also allows to (re-)examine the size of the attended area, which had yielded divergent results (e.g., 59-62). Moreover, it can shed new light on a fundamental question of attention research, namely whether attention is *space-based* or *object-based* (e.g., 63-66; for a review, see 67). Finally, accurate estimation of the size and shape of the attention focus, as well as its temporal and spatial dynamics during voluntary (endogenous) or automatic (exogenous) attentional orienting, or prior to goal-directed actions has crucial implications for conceptual and computational models of visual attention (e.g., 7,50,68-70).

Objects undeniably define the scene (71,72) and guide our actions and memories (73-75). For this reason, there are numerous scientific questions for which an object-based measurement is preferable. Several other questions, however, are better investigated in the absence of localized test items. For instance, when evaluating the distribution of attention surrounding salient or relevant objects (*exogenous* and *endogenous attention*) or when assessing the spread of attention across visual structures (*object-based attention*) or between presented items (e.g., during *visual search*). Moreover, the item-free 1/f noise protocol is particularly suitable to study attentional modulations in the context of motor actions (*premotor attention –* for a recent proof-of-concept see 66) and offers promising *clinical applications*, as it for instance allows to precisely map out regions with perceptual or attentional deficits.

To conclude, the presented protocol offers several decisive advantages over conventional paradigms that rely on transient, localized test stimuli to measure the perceptual correlates of visual attention. By neither conveying spatial nor temporal information on where or when the test signal is presented, the stimulus does not bias observers’ expectations and thus enables an undistorted attention assessment. Since visual sensitivity for local 1/f orientation signals does not fall off with retinal eccentricity, the protocol allows to directly compare performance from fovea to periphery, without the requirement to scale the signal strength with retinal distance. These unique features render the dynamic noise protocol an ideal tool to map the distribution of visual attention across space and time – and to address research questions that could not yet, or only to a limited extent, be investigated with conventional, object-based approaches.

## Declarations

### Funding

This research was supported by grants of the Deutsche Forschungsgemeinschaft (DFG) to HD (DE336/5-1) and a Marie Skłodowska-Curie individual fellowship by the European Commission to NMH (898520).

### Competing interests

The authors declare no competing interests.

### Ethics Approval

The protocols for the study were approved by the ethical review board of the Faculty of Psychology and Education of the Ludwig-Maximilians-Universität München (approval number 13_b_2015), in accordance with German regulations and the Declaration of Helsinki.

### Consent to participate

All observers gave written informed consent.

### Author contributions

Conceptualization and methodology: NMH, HD; Software: NMH; Investigation: NMH; Formal analysis: NMH; Visualization: NMH; Writing—original draft: NMH; Writing—review & editing: NMH, HD; Funding acquisition: NMH, HD.

## Acknowledgements

We are grateful to the members of the Deubel lab, in particular to David Aagten-Murphy, and to Dejan Draschkow for helpful comments and discussions.

## Open Practices Statement

Eyetracking and behavioral data will be publicly available at OSF upon manuscript publication, custom MATLAB code for generating the 1/f stimulus will be publicly available on GitHub. None of the experiments was preregistered.

## Additional information and files

**Supplemental Video S1**

Demonstration of stimuli and design of Experiment 1a (*cued trial*).

**Supplemental Video S2**

Demonstration of stimuli and design of Experiment 1a (*uncued trial*).

## Notes

### Competing Interest Statement

The authors have declared no competing interest.

### Summary of Updates

Section order updated; Supplemental files updated.

